# Feasibility of improving manufacturability based on protein engineering

**DOI:** 10.1101/2023.11.23.568267

**Authors:** Florian Capito, Ting Hin Wong, Christine Faust, Kilian Brand, Werner Dittrich, Mark Sommerfeld, Thomas Langer

## Abstract

While bioactivity and a favorable safety profile for biotherapeutics is of utmost importance, manufacturability is also worth of consideration to ease the manufacturing process. Many biotherapeutics are typically expressed in mammalian cells. Process-related impurities or biological impurities like viruses and host cell proteins (HCP) are present in the harvest which have mostly acid isoelectric points and need to be removed to ensure safety for the patients. Therefore, during molecule design, an isoelectric point of the target molecule should preferably differ sufficiently from the isoelectric points of the impurities to enable an efficient and straightforward purification strategy. In this feasibility study we have evaluated the possibility to improve manufacturability by increasing the isoelectric point of the target protein. We have generated several variants of a GLP1-receptor-agonist-Fc-domain -FGF21 fusion protein and demonstrate that the critical anion exchange chromatography step can be run at high pH values with maximal product recovery theoretically allowing removal of HCP and viruses. Addressing the isoelectric point can be useful for an efficient process for removing HCP and viruses and this topic should be considered early in the research phase to ensure that other important molecule properties, e.g. safety, efficacy and expression yield are not impacted.

## 1. Introduction

Monoclonal antibodies (mAbs) have become the most promising molecules for medical care over the past years and the number of molecules reaching the clinical stage is still increasing year by year. As of November 2022 more than 1200 clinical trials for antibody therapeutics are ongoing (Kaplon et al., 2022). Most therapeutic mAbs are typically being purified using protein A chromatography followed by 1-2 polishing steps. These polishing steps may comprise a cation-exchange (CEX) chromatography step, an anion-exchange (AEX)-chromatography, mixed-mode chromatography or a hydrophobic interaction chromatography (Shukla et al., 2007). As most antibodies have basic isoelectric points, an AEX-chromatography step is the first choice for a polishing step. This step is operated in flow-through mode with the target protein passing through the resin while impurities are bound to the resin. In addition, the AEX-chromatography step is important for virus removal. Since most impurities, such as viruses, e.g. the murine leukemia virus (MuLV) and minute virus of mice (MVM) as well as host cell proteins have acidic isoelectric points (Qasim et al., 2011) the implementation of an AEX-chromatography step is advantageous with regards to yields and costs. Hence, for mAbs platform-processes emerged based on the implementation of a protein A capture step followed by an AEX-chromatography step (Shukla et al., 2017). Yet, with the emergence of novel protein formats, e.g. Fc-fusion proteins, the typical platform process for mAbs may not be applicable any longer and, depending on the physico-chemical properties of these molecules, tailor-made purification processes need to be established. This is acceptable as finding a molecule with desired bioactivity and efficacy profile while at the same time ensuring a favorable safety profile may be hard enough. Yet, if safety and efficacy are not impacted, it may be worth to invest in additional protein engineering rounds addressing manufacturability to avoid longer development times and overall higher costs. In the past, developability assessments were mostly focused on formulate-ability and long-term stability to asses oxidation, degradation, and aggregation of the molecules and to a lesser extent on manufacturability (Yang et al., 2013; Jarasch et al., 2015). *In silico* models to evaluate interactions with excipients depending on peptide physico-chemical properties are also considered during drug development. A thorough developability assessment prior transfer of a drug candidate into development can be beneficial as it may avoid severe manufacturing and stability risks which otherwise may require a re-design of the drug-candidate (Shan et al., 2018; Evers et al., 2018). In order to keep the time-to-clinic as short as possible it is crucial to address all aspects of developability, i.e. manufacturability as well as formulate-ability as early as possible during the research phase. This means that the impact of the physico-chemical properties of a drug candidate should not be disregarded as these properties will determine whether and how fast a process for a drug candidate can be developed, as long as changes in physico-chemical properties have no adverse effects on bioactivity and safety of the drug. As corroborated by others one should also take into account to adjust the physico-chemical properties of a drug candidate during the developability studies. This adaption can in principal be done without losing biological activity (Bak & Dai, 2015; Bak et al., 2015; Fosgerau & Hoffmann, 2015; Shan et al., 2018).

Manufacturability describes „how easy a molecule can be manufactured“, *i.e.* expressed and purified. Conley et al. describe a manufacturability design to optimize the amino acid sequence with regards to protein aggregation. Due to unfavorable physico-chemical properties of their original mAb, a straightforward purification process was not possible. A new purification process needed to be set up. This included an alternative non-protein A capture step with challenges on impurity removal, safety requirements and additional studies for optimization and process validation. By addressing the observed manufacturability issues and starting a protein engineering approach a new mAb variant was created without impact safety and efficacy. This re-designed antibody was fit for platform and finally used in pre-clinical studies (Conley et al., 2011).

The present work deals with a fusion protein containing a N-terminal glucagon-like peptide 1-receptor-agonist (GLP1-RA) sequence, an fragment crystallizable (Fc)-domain from human immunoglobulin, and a FGF21 variant. GLP1 belongs to the incretins which are released postprandial by the gut. The main function of GLP1 is to stimulate insulin secretion. This so-called incretin-effect is almost absent in patients with type 2 diabetes. However, exogenously administered GLP1-RA is able to elicit insulin secretion and hence able to normalize blood plasma glucose levels (Andersen et al., 2018). FGF21 was first cloned in 2000 and was placed due to its sequence into the FGF family (Nishimura et al., 2000). While conventional members of the FGF family act in a paracrine or autocrine manner, FGF21 belongs to an atypical FGF subfamily that can enter the circulation and hence acts as an endocrine hormone. The extensive and pleiotropic pharmacologic effects of FGF21 includes weight loss, improvement of insulin sensitivity without causing hypoglycemia, and significant improvements of plasma triglyceride and cholesterol levels (Coskun et al., 2008; Kharitonenkov et al., 2005; Xu et al., 2009; Adams et al., 2013; Kharitonenkov et al., 2007; Talukdar et al., 2016; Veniant et al., 2012). This metabolic profile makes FGF21 a potential diabetes and obesity drug and in addition also favorable for the treatment of non-alcoholic fatty liver disease and its more severe stage, steatohepatitis (NASH). By now several FGF21-derivatives have already entered the clinical stage (Chen et al., 2022). Furthermore, it was found that the combination of FGF21 and a GLP1-RA has synergistic beneficial effects (Hong et al., 2016) and corresponding fusion proteins have been designed. Such constructs also addressed the need for an enhanced plasma half-life, since GLP1-RA and FGF21 suffer from short plasma half-lives (Pan et al., 2021; Ye et al., 2023). Using the Fc-domain as fusion partner is a well-established method to enhance the plasma half-live of biomolecules (Liu, 2018).

The present work describes a feasibility study to assess the possibilities to improve manufacturability in downstream processing by applying protein engineering to the protein of interest. The calculated isoelectric points of the initial protein variants are 6.15 and 6.62. An isoelectric point of > 7 is considered to be beneficial for employing an efficient AEX-chromatography step, where the target protein is in the flow through while host cell proteins and viruses bind to the AEX-matrix. Therefore we have made several variants of GLP1-RA-Fc-FGF21 fusion proteins with altered isoelectric points in order to evaluate the impact of the isoelectric point on the yield in the AEX-chromatography step.

## 2. Materials and Methods

### 2.1. Protein constructs and protein expression

Two sets of fusion proteins with two different GLP1-RAs, designated variant A and B, were created. To alter the isoelectric point, different numbers of lysine-residues were either inserted into the linker sequence between the GLP1-RA and the Fc-domain or attached to the C-terminus of the protein. Additional constructs with lower isoelectric points were made by adding a poly-glutamate stretch at the C-terminus. The DNA coding for the protein sequences was synthesized using codon-optimization for human host cells (Thermo Fisher Scientific). If the codon optimization for the lysine-stretches resulted in six or more consecutive adenines in the DNA, a second gene was synthesized. In these genes all lysines in the linker sequences are encoded by the AAG codon, whereas the rest of the genes remained unchanged with regard to codon usage. The isoelectric points of the proteins were calculated using an in-house tool based on the EMBOSS-software (Rice et al., 2000).

All protein variants were cloned into an expression vector under a CMV promotor and a leader sequence directing the proteins into the culture supernatant. Protein expression was achieved by transient transfection of FreeStyle human embryonic kidney 293 (HEK293-F) cells (Thermo Fisher Scientific). Cells were grown in non-baffled shake flasks (Corning) at 110 rpm, 37°C and 8% CO_2_. Cells used for transfection were grown to a cell density of approx. 1.2 × 10^6^ cells/mL. For transfection, DNA was mixed with linear polyethyleneimine (PEI) at a ratio of 1 : 3 in Opti-MEM I-medium (Thermo Fisher Scientific). The transfection mixture was incubated for 20 minutes at room temperature and then added to the cell cultures. Cells were cultivated using Freestyle F17 medium supplemented with 6 mmol/L glutamine. After 6 days the culture supernatant was separated from the cells by centrifugation (30 min at 4.500 × *g* and 4°C). Cell pellets were discarded and the supernatants were cleared by 0.22 µm sterile filtration. Cleared culture supernatants were used for further work. The titer in the cleared culture was determined by biolayer interferometry using the Blitz-system (Pall Fortebio, Fremont, CA, USA). FGF21 was used as a control protein in the activity assay. Expression of human full length FGF21 protein was done in *E.coli.* Protein was recovered from inclusion bodies following a procedure described by Hecht et al. (2012). Additionally, the initial protein variants A and B were expressed in chinese hamster ovary (CHO) cells. This was done using stable transfected cell pools using proprietary cell media and cultivation conditions. The proteins obtained from CHO cells were used for optimizing the AEX-chromatography step conditions (different load ratio under different pH and conductivity).

### 2.2. Purification of fusion proteins

For all protein variants a protein A chromatography (MabSelect SuRe resin, GE Healthcare, Uppsala, Sweden) was used as capture step. Further purification was done in 96 well microplates using Capto Q AEX-resin (GE Healthcare).

The pH and conductivity were varied in the 96 well microtiterplate for the CHO derived variants, with prior equilibration of the protein to be loaded on the AEX to the corresponding pH and conductivity values. In addition, the load ratio was varied for the CHO derived variant. For variants expressed in HEK293 cells, due to limited availability of material, only pH 6.0 and pH 8.0 conditions were compared in head-to-head experiments.

Purification step yields of 96 well microtiterplates was determined by UV280nm measurements using a Lunatic system (Unchained Labs, Pleasontan, CA, USA). HCP levels in the AEX flow-through of the CHO derived variant were determined by 3^rd^ generation HCP-ELISA kit (Cygnus Technologies, Southport, NC, USA). For HEK 293 derived variants, HCP-levels were not quantified due to limited available material. Contour plots showing the HCP levels as well as obtained purification yields were generated using the SigmaPlot software (version 14, Systat Software GmbH, Erkrath, Germany).

### 2.3. SDS-PAGE analysis and isoelectric focusing

Protein samples (5 µg protein) were mixed with either 4 x LDS sample buffer (Thermo-Fisher Life Technologies) or 4x LDS sample buffer + 50 mM dithiothreitol for SDS-PAGE analysis under non-reducing or reducing conditions, respectively. Samples were incubated for 5 min at 99°C before loading on 4-12 % SDS-PAGE with MOPS as running buffer (Thermo-Fisher Life Technologies). BenchMark protein ladder was used as marker (Invitrogen). Isoelectric focusing was done using IEF gels ServaGel IEF 3-10 (Serva, Heidelberg, Germany) using the corresponding anode and cathode buffers (Serva). 15 µg protein samples were mixed with IEF pH 3-10 sample buffer (Serva) and loaded immediately on the gels. Marker proteins were from the isoelectric focusing calibration kit (high pH range, GE Healthcare).

### 2.4. Measurement of biological activity

Biological activity of selected variants was tested using a luciferase bioluminescence assay. Briefly, the reporter cell line iLite FGF21 Assay Ready Cells (Svar Life Science AB formerly EuroDiagnostika, Malmö, Sweden) overexpressing the human FGFR1c together with β-klotho was plated into 384-well micro-titer plates (Perkin Elmer, Cat. #6007480) and incubated at 37°C, 5% CO_2_ and 95% relative humidity. For dose-response curves each fusion protein was 5-fold diluted in quadruplicates starting at 100 nmol/L for a 12-point curve, added to the cells and incubated for 5 h. Reagents used to analyze Luciferase activity were from the Dual-Glo Luciferase Assay System (Promega, E2940) according to the supplier’s protocol. Luminescence was read using a multi-mode plate reader (CLARIOstar, BMG Labtech, Germany).

## 3. Results and Discussion

### 3.1. Protein expression

The schematic structures of the protein variants are shown in Fig. 1. To alter the isoelectric point charged amino acids residues were either introduced into the linker sequence or attached at the C-terminus of the protein. All protein variants were expressed in mammalian cells and the protein titer was measured in the supernatant using biolayer interferometry. An unexpected result was that the expression of the protein variants containing the introduced lysine-residues in the linker sequence was very low. In contrast to this, the expression level of protein variant A-C(E6) was roughly double as high as the expression level of the initial variant A. Expression level of variant A-C(K6) was in the same range as of the initial variant A. For the protein variant B we observed no enhanced expression upon fusing charged amino acids to the C-terminus. The expression levels were in the range of the initial variant B. For the very low expressing constructs, we observed no unusual incidents during cell cultivation with regards to cell viability and cell density. However, an impact of the expression levels of proteins as a function of consecutive adenines in the coding DNA was described earlier. In these reports reduced expression levels and the occurrence of truncated protein variants was attributed to premature translation stop and ribosome sliding (Arthur et al., 2015; Koutmou et al., 2015). To check whether the low expression levels was due to the presence of six or more consecutive lysine residues we created a second set of protein variants. In these constructs the AAA-codons for lysine-residues in the linker sequences were replaced by the alternative AAG-codon. The protein and DNA-sequences for the corresponding regions are depicted in Fig. 2. However, the expression levels for the variants A with 8 and 15 lysines containing the AAG codons are slightly enhanced, but still very low compared to the initial variant. The expression level for the variant A-L(K15AAG) is even lower than the expression level for the corresponding variant with the AAA-codons for lysines. For variant B the replacement of AAA codons with AAG codons did not alter the expression levels at all. The expression levels for the different protein variants are shown in Fig. 3.

**Figure 1.**
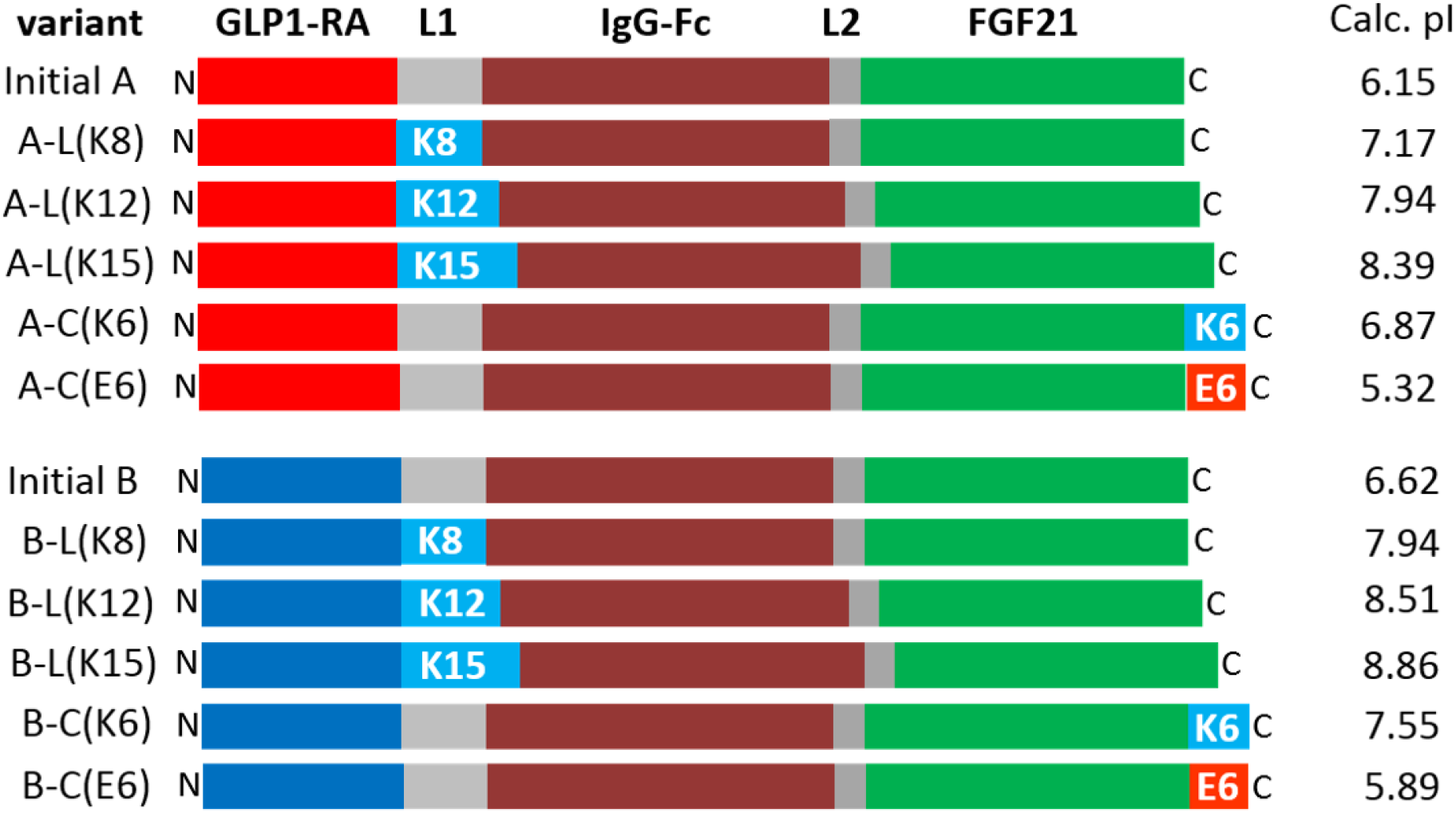
Schematic depiction of the various protein variants. GLP1-RA is an activating ligand for the GLP1-receptor. Two GLP1-RA variants, designated variant A and B, were used. L1: linker sequence between GLP1-RA and the Fc-domain. L2: linker sequence between the Fc-domain and FGF21. The L2 sequence is unchanged throughout all constructs.

**Figure 2.**
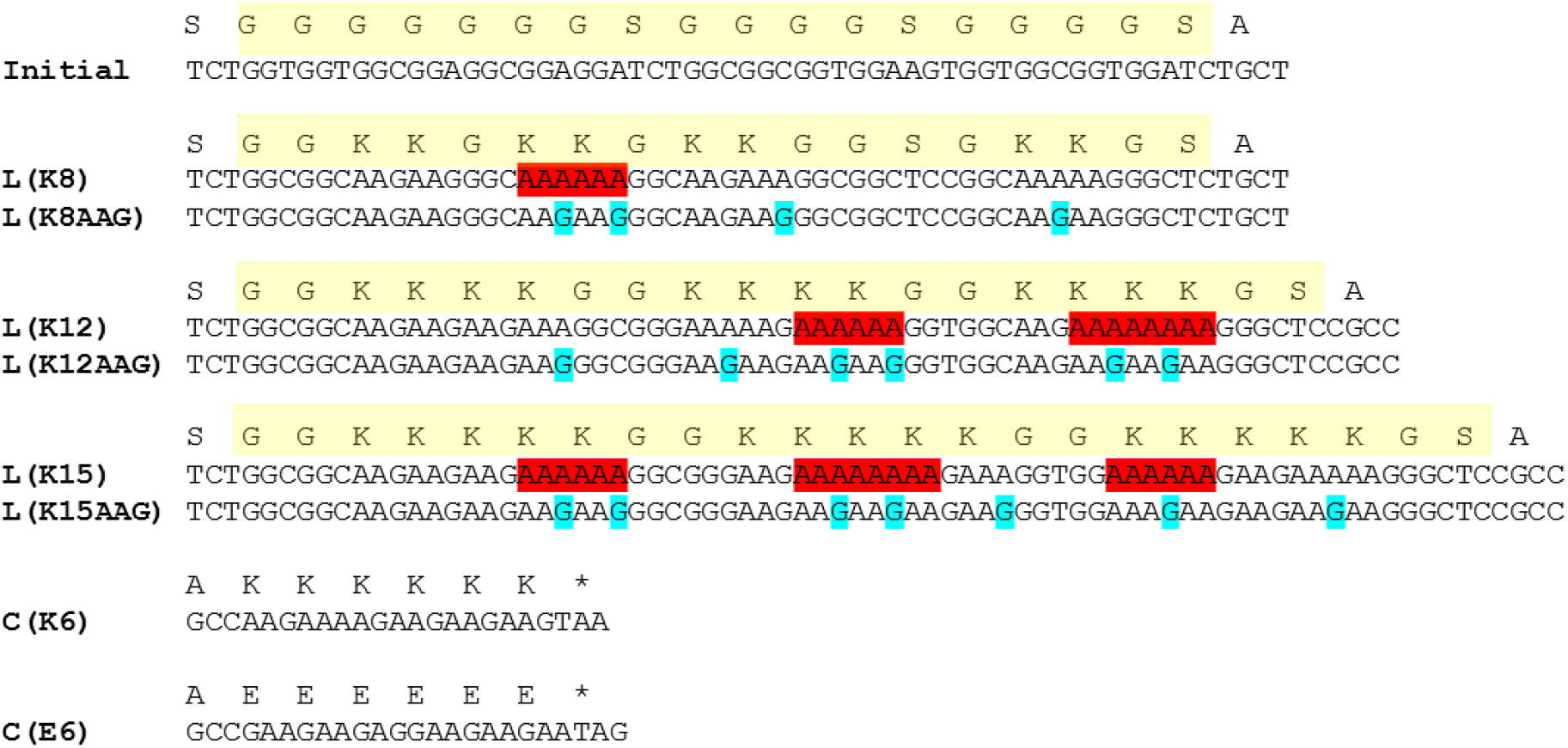
Sequences of the different L1-linker and the C-terminal charged tags. The protein linker sequence is highlighted in yellow. Stretches of more than six consecutive adenines are highlighted in red. Changes for the adapted constructs are marked in cyan. In the adapted constructs the AAG codon was consequently used for all lysines.

**Figure 3.**
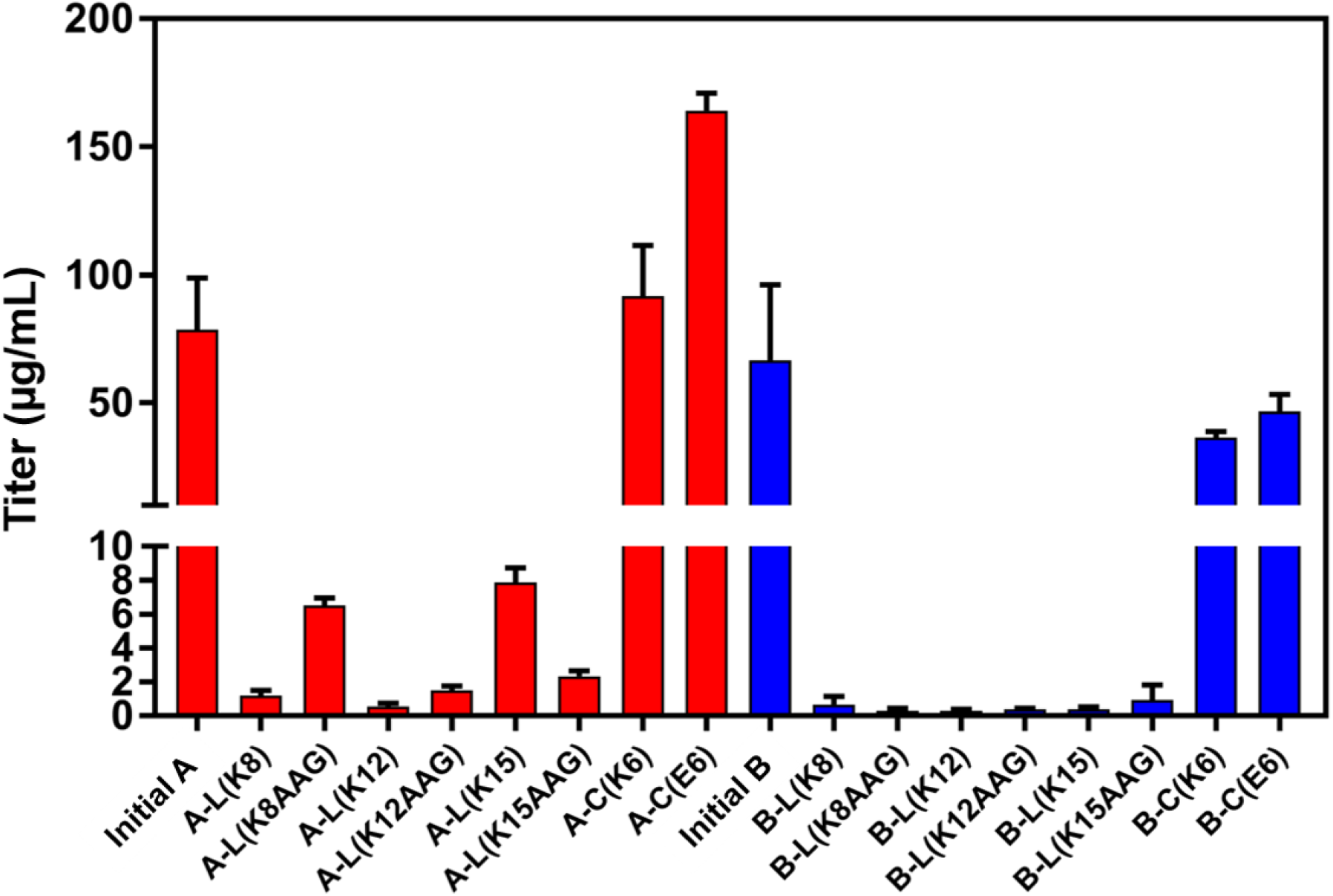
Expression levels of the different protein variants as measured in the cleared supernatants. Remarkable is the loss of expression level for the variants with lysine-residues inserted in the linker sequences, even after replacement of the AAA codons with AAG codons.

Arthur et al. checked the expression of a mCherry reporter protein that was placed behind different poly-lysine-stretches with western blot and fluorescence assays as well as on the mRNA level. With six consecutive AAA codons (18 consecutive adenines) they observed a drop of approximately 3 fold in protein expression. This reduction is correlated with a drop in the amount of detected mRNA. When analyzing variants containing consecutive AAG codons they observed nearly the same expression levels for protein and mRNA for six AAG codons, and only a slight drop in expression when nine AAG codons are placed before the reporter protein (Arthur et al., 2015). However, in our constructs maximal eight consecutive adenines were present (Fig 2). But even after replacing the AAA codons with AAG codons the reduction in protein expression could not be rescued. The reason for the low expression levels that we have observed remains still obscure. Obviously, other factors than the presence of poly-adenine-stretches are important for proper protein expression.

As protein expression levels were higher for the A-variants we have only analyzed these variants by SDS-PAGE and isoelectric focusing. Due to the presence of the Fc-domain the proteins are dimeric, as long as the connecting disulfide-bonds are not reduced. The calculated molecular weights for our proteins of interest are ∼51 kDa (monomer, reduced) and ∼102 kDa (dimer, non-reduced). The observed apparent molecular weights are within this range (Fig. 4a). Results from the isoelectric focusing reveal some heterogeneity of the analyzed protein samples (Fig. 4b). N-glycosylation is a common modification of antibodies which occurs at a conserved Asn residue in the CH2 part of the Fc-domain. This is also true for Fc-fusion molecules (Liu, 2018). N-glycosylation can be very heterogeneous and differences in the content of sialic acid lead to differently charged isoforms (Croset et al., 2012; Beck & Liu, 2019). However, the obtained isoelectric point for the initial variant A is ∼6.0, which is close to the calculated isoelectric point. The calculated isoelectric point for the variant A-C(E6) was 5.32. However, the obtained isoelectric of ∼5.8 point is slightly higher. For the variant A-C(K6) the main band is at the same height as that of the initial variant A. A faint “ladder” above the main band is visible. This might be indicative for a C-terminal successive clipping of the lysine-residues. The calculated isoelectric point for variant A-L(K8) was 7.17. In this case the observed isoelectric point (∼7.5) is slightly higher than the calculated isoelectric point. For the protein variants A-L(K12) and A-L(K15) we could not determine the isoelectric points with our experimental setup propably because the isoelectric points are > 8.3 and thus the proteins do not migrate into the gel (data not shown).

**Figure 4.**
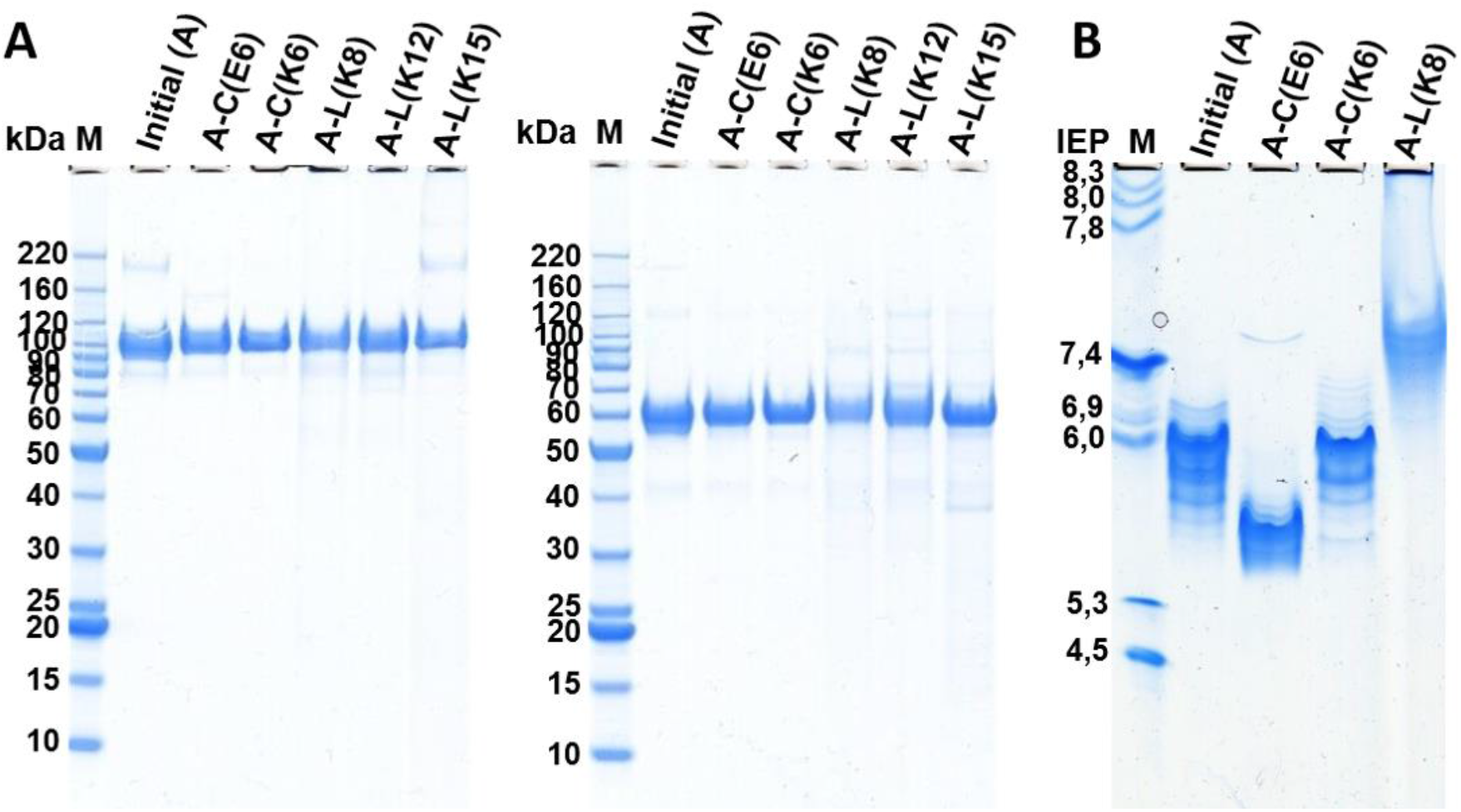
SDS-PAGE under non-reducing and reducing conditions (A) and isoelectric focusing (B) of the initial variant A and altered versions of hereof. Expression was done in HEK 293 cells. The label of the protein does not consider which expression vector was used. Protein samples used were from elution of the protein A column.

### 3.3. Impurity removal and anion exchange chromatography step yield – need to raise the isoelectric point

The initial variants A and B were produced from a stable transfected cell line pool. We focused on the variant B, which has a higher isoelectric point compared to variant A (6.62 compared to 6.15). For a protein intended to be used as a biotherapeutic an effective purification strategy is as well elementary as the removal of HCP (Tuameh et al. 2022). We have analyzed the HCP content in the flow-through of the AEX-chromatography step as a function of pH and load ratio of the column. Due to the rather low isoelectric points of most HCP, the HCP level in the flow-through of the AEX-chromatography is usually lower when the AEX-chromatography is performed at elevated pH values. This has also been corroborated by others showing better HCP clearance when conducting the AEX-chromatography at elevated pH conditions with significant higher log reduction for HCP at pH 8.5 and 9.0 compared to lower pH values (Kelley et al., 2008; Weaver et al., 2013). In addition, the load ratio has an impact on the HCP level: At high load ratios the binding sites on the resin are already occupied with HCP and thus can not bind more impurities. This results in elevated HCP levels. Contrary, at low load ratios sufficient binding sites on the resin are available to remove these impurities. However, as the isoelectric point of the analyzed protein variant is rather low, the yield for the purification step shows an opposite correlation. The corresponding contour plots are shown in Fig. 5. The yield for the purification step is as low as 30-60% -depending on the load ratio-when the purification step is performed at pH 7.3. The maximum yield for this purification step could be raised to approximately 70% – but for the sake of elevated HCP levels. Other points to consider are the typical model viruses (*e.g.* MuLV, MVM) used during virus clearance studies prior producing material for clinical trials. These viruses have isoelectric points in the range of ∼5 – 6 (Anouja, et al., 1997; Strauss et al., 2009a). In addition, most animal viruses also have isoelectric points in the slightly acidic range (Michen & Graule, 2010; Miesegaes et al. 2010). Hence it is desirable to run the AEX-chromatography step at high pH (∼8 – 8.5) to ensure a robust virus clearance. Just as for HCP removal virus clearance is more efficient at higher pH values.

**Figure 5.**
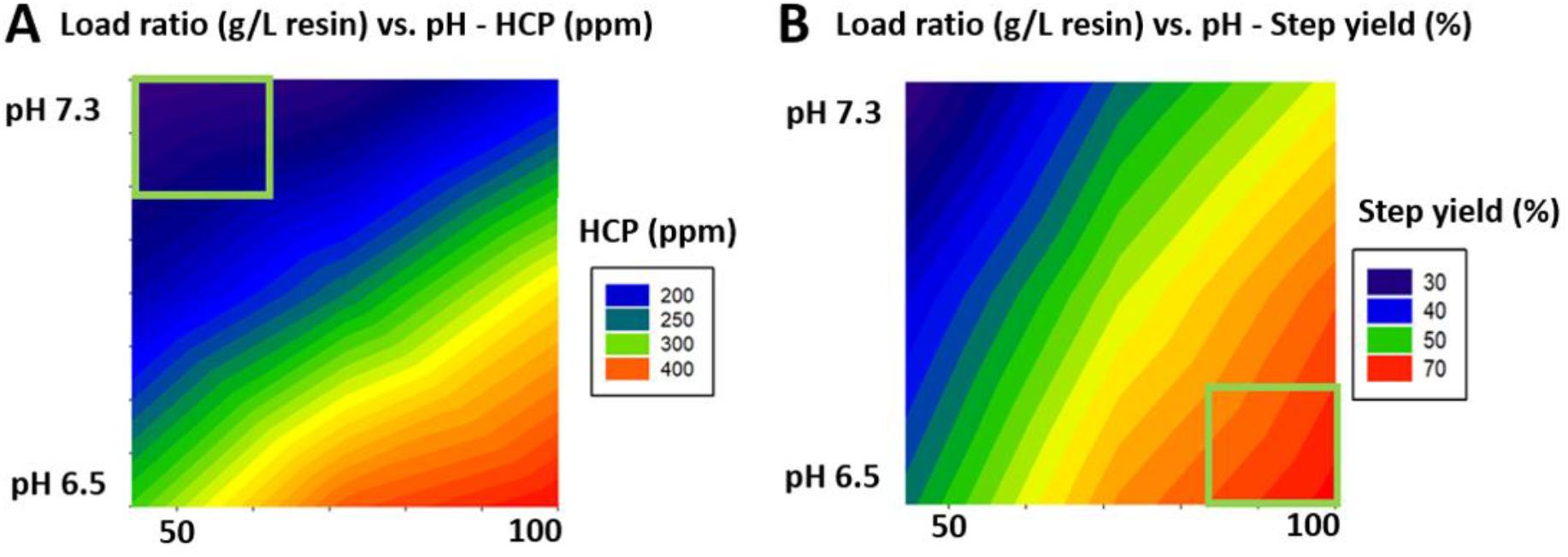
Contour plots showing the dependence of (A) HCP clearance and (B) the yield in the AEX-chromatography step as a function of pH and the load ratio (g target protein / L resin). For these experiments, initial variant B (isolectric point 6.62) was used. The HCP levels were measured in the flow-through from the AEX-chromatography step. The step yield is given as the percentage of the recovered protein from the AEX-chromatography step. The preferred operating conditions for the corresponding issues are marked with a green box. An elevated pH value is preferred for HCP removal whereas a lower pH is beneficial for the step yield.

Derived from this choosing a protein with a rather low isoelectric point will require additional work to ensure robust virus clearance while maintaining high yield in downstream processing. At the end, this will require the replacement of the AEX-chromatography step by an alternative purification step(s) with the consequence to establish and evaluate the virus and HCP removal process. An attractive alternative to establish and validate a new purification step is to alter the isoelectric point of the protein of interest. We have analyzed the step yield for the AEX-chromatography step for different protein variants with different isoelectric points at pH 6.0 and pH 8.0 (Fig. 6). The AEX-chromatography step yield for the initial protein variants A and B at a load rate of 40 g/L was >20% at pH 6.0 and <5% at pH 8.0. For protein variants with isoelectric points ∼8.0 and higher the AEX-chromatography step yield could be raised to > 90% at pH 6.0 as well as up to 85% at pH 8.0. Based on findings by Curtis and coworkers (Curtis et al. 2003), showing better clearance of simian virus at pH 8.0 compared to lower pH with an AEX-chromatography step, this would also improve virus clearance. They also pointed to the fact that target proteins with a similar isoelectric point compared to the isoelectric points of the viruses may make the AEX-chromatography step inefficient regarding to virus removal. Previous studies showed that interaction between viruses and AEX-chromatography resin follows primarily electrostatic interactions. Thus, based on the isoelectric points of the viruses, enhanced binding to the AEX-chromatography-resin is expected when raising the pH above their isoelectric points, although this is also dependent on the virus species and charge distribution on its surface (Strauss et al. 2009a,b). Thus, running the AEX-chromatography step at higher pH values would generally allow for improvement of virus clearance, considering the typically lower isoelectric points of viruses occurring in mammalian cell cultures. As expression levels for most protein variants were surprisingly low, HCP clearance with the AEX-chromatography step was only evaluated for one variant. Yet, obtained results (see Fig. 5) shows a general tendency of improved clearance at elevated pH values. This is in accordance with results from Kelley and coworkers demonstrating improved HCP clearance when running the AEX-chromatography step at elevated pH values (Kelley et al. 2008).

**Figure 6.**
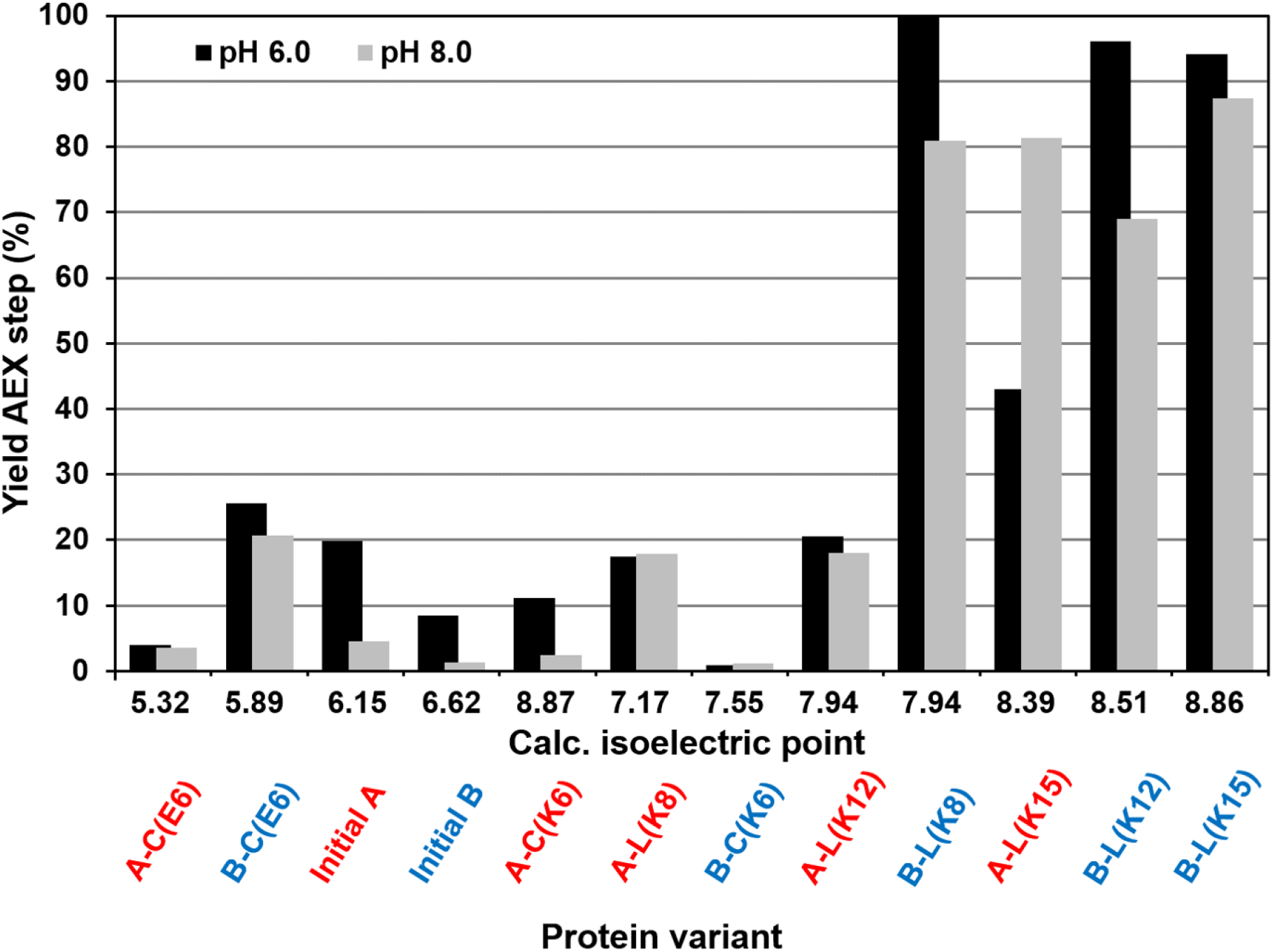
Impact of the isoelectric point of the protein of interest on the yield for AEX-chromatography. The different protein variants were arranged according their calculated isoelectric points. The AEX-chromatography step was either done at pH 6.0 (black bars) or pH 8.0 (grey bars). Note: Since the protein expression was very poor for the variants A-L(K8), A-L(K12), B-L(K8), A-L(K15) and B-L(K12) the load ratio for these variants was only between 13 and 25 g/L (compared to 40 g/L for all other variants). Yet, achieved step yields were still high for some of these variants, despite underloading the resin.

We have noticed that the step yields of the protein variants with lysine-residues at the C-terminus do not differ markedly from the initial variants. This finding is in accordance to the results from the isoelectric focusing: The C-terminal lysine-residues obviously have been cleaved off during expression. While not being the case in this feasibility approach with the given target molecule and feedstock parameters, for some circumstances, it may also be envisioned, to allow the implementation of a two-column process using target molecules with raised isoelectric point. Yet, this requires very low HCP levels already in the feedstock and robust removal to below drug substance specifications with such a process. Finally, an increased isoelectric point of the protein of interest is likely to result in an enlarged “sweet spot” (s. Fig.5) – the realization of a higher AEX-chromatography step yield as well as good impurity depletion.

### 3.5. Impact on biological activity

The biological activity of FGF21 and selected GLP1-RA-Fc-FGF21 fusion proteins was tested in a luciferase bioluminescence assay (Fig. 7). Fusion of FGF21 to a GLP1-RA as well as to the Fc-domain might have some impact on the activity of the FGF21 protein. The initial variant A had a reduced EC50 value compared to FGF21 and the initial variant B. However, maximal receptor response could be evoked for this construct. No activity could be measured for the protein variants with addition of glutamate-residues to the C-terminus. This means that the additional amino acids or charges prevent the correct binding to the receptor and hence its activation. The structure of the C-terminus from FGF21 in complex with the receptor β-klotho has been elucidated recently (Lee et al., 2018). The structure reveals that the C-terminal amino acids of FGF21 are bound in a deep cleft and also the last amino acid of the C-terminus is buried into the substrate binding pocket of β-klotho. This binding modus does obviously not allow a C-terminal extension of FGF21. The protein variants with additional lysine-residues at the C-terminus showed similar activity as the initial variants. These findings are in accordance with the observation that the C-terminal lysine-residues have been clipped (see Fig. 4b). Clipping of the C-terminal lysine is widely observed when full length mAbs are expressed by CHO-cell cultures – unless the CHO cells have been genetically engineered to knock out the corresponding protease (Gramer, 2014; Hu et al., 2016). Activity for protein variants with internal lysine-residues A-L(K8), A-L(K12) and A-L(K15) have also been measured (Fig. 7b). The activity of these variants differto some extent from the original variant. As noted above, the FGF21 part of the molecule is unchanged and also the initial protein variants A and B exhibit different activities in the FGF21 activity assay. Note, that the included lysine-residues are spatial distant from the FGF21 domain (see Fig. 1). It is unknown in which manner these additional amino acids, either the different GLP1-RA-variants or the internal lysine-residues, leads to the observed difference in the activity assay. It has yet to be evaluated how such modifications translate into *in vivo*, i.e. have impact on the pharmacodynamic and pharmacokinetic properties of the protein.

**Figure 7.**
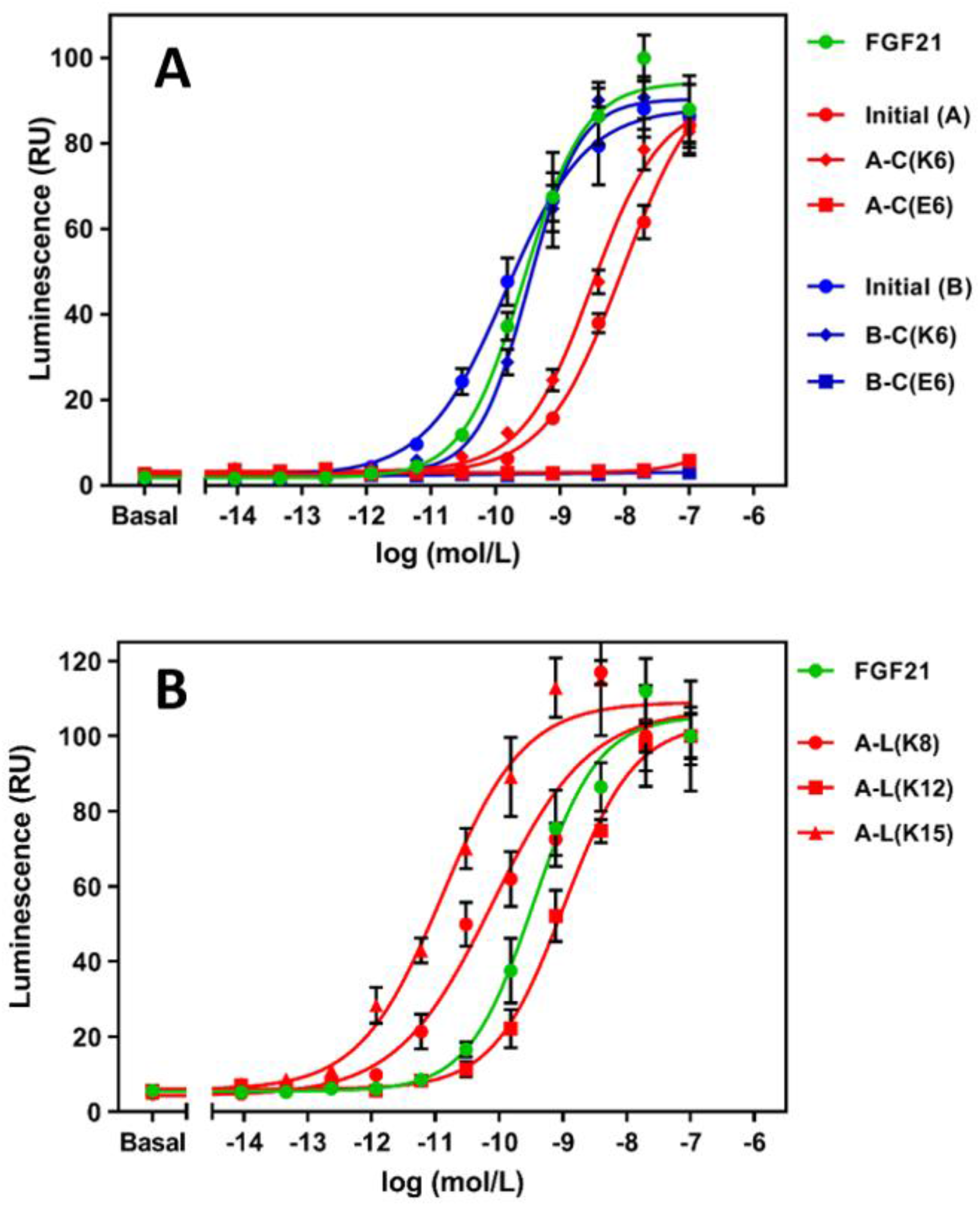
Dose-response curves from a luciferase gene expression assay with HEK293 cells overexpressing the human FGFR1 and β-klotho proteins showing the biological activity of selected variants. Measured luminescence (in relative units) as a function of protein concentration as determined with the luciferase activity assay. The average of four biological replicas ± SEM is shown.

## 4. Conclusions

While efficacy and safety are almost or primary concern when developing a drug candidate, the importance of considering the isoelectric point for a potential therapeutic protein was evaluated in this feasibility study. A significant increase in step yield for the AEX-chromatography and an improved overall purification yield can be achieved when using protein variants with enhanced isoelectric points compared to the original low isoelectric point variants. This might allow the usage of mAb purification platforms and, depending on impurity levels, potentially two-column processes and hence shortening of the development timelines and development efforts. In addition, better virus clearance and impurity removal (HCP) are anticipated based also on earlier work and the mechanism of virus and HCP binding to the anion-exchange-chromatography resin (Curtis et al, 2003; Kelley et al. 2008). Furthermore, a more basic isoelectric point might also lead to an increased solubility when moving the isoelectric point away from the pH of the formulation buffer (Tan et al., 1998).

Our main intend with this feasibility study was to pinpoint on the importance of considering the isoelectric point of a potential therapeutic protein for manufacturability, secondary to efficacy and safety. Introducing of additional charged amino acids or mutating amino acids in a protein sequence can have fundamental impact on other protein properties. In our case, the most dramatic impact of such a kind of protein engineering on the protein properties has been observed by the introduction of lysine-rich stretches in the linker-sequence between the GLP1-RA and the Fc-domain. This resulted in a dramatic loss of protein expression yields addressing now a different point for the manufacturability of potential therapeutic proteins. In addition slightly changed biological activities of these variants were observed. Additional studies are needed to make a statement whether this has an impact on the *in vivo* function of the molecule or whether yet not anticipated properties are affected, e.g. enhanced adverse effects or other safety issues. Also, we observed C-terminal clipping of the C-terminal added lysine-residues which resulted in the production of proteins comparable to the initial variants.

Of course, maintaining the biological function is crucial. With our feasibility study we want to emphasize the importance of addressing the manufacturability and the importance of the isoelectric point not only during development but already during the discovery phase. This is of utmost importance for a fast-to-clinic track. Otherwise, certain drawbacks (e.g. need for cell line engineering, implementation of non-platform processes, coping with suboptimal process steps – which are all time-consuming procedures) are unavoidable.

## Acknowledgements

Conflict of interest statement: C.F., K.B., W.D., M.S, T.L. and F.C. are employees of Sanofi-Aventis Deutschland GmbH and may hold company shares and/or stock options. T.H.W. was during the time of work a registered student at the THM Giessen, 35390 Gießen, and working at Sanofi-Aventis Deutschland GmbH and may hold company shares and/or stock options.

